# A comprehensive mouse kidney atlas enables rare cell population characterization and robust marker discovery

**DOI:** 10.1101/2022.07.02.498501

**Authors:** Claudio Novella-Rausell, Magda Grudniewska, Dorien J. M. Peters, Ahmed Mahfouz

## Abstract

The cellular diversity and complexity of the kidney are on par with its physiological intricacy. Although our anatomical understanding of the different segments and their functions is supported by a plethora of research, the identification of distinct and rare cell populations and their markers remains elusive. Here, we leverage the large number of cells and nuclei profiles using single-cell (scRNA-seq) and single-nuclei (snRNA-seq) RNA-sequencing to build a comprehensive atlas of the adult mouse kidney. We created MKA (<u>M</u>ouse <u>K</u>idney <u>A</u>tlas) by integrating 59 publicly available single-cell and single-nuclei transcriptomic datasets from eight independent studies. The atlas contains more than 140.000 cells and nuclei covering different single-cell technologies, age, and tissue sections. To harmonize annotations across datasets, we constructed a hierarchical model of the cell populations present in our atlas. Using this hierarchy, we trained a model to automatically identify cells in unannotated datasets and evaluated its performance against well-established methods and annotation references. Our learnt model is dynamic, allowing the incorporation of novel cell populations and refinement of known profiles as more datasets become available. Using MKA and the learned model of cellular hierarchies, we predicted previously missing cell annotations from several studies and characterized well-studied and rare cell populations. This allowed us to identify reproducible markers across studies for poorly understood cell types and transitional states.

## Introduction

Kidneys are organs with a high degree of cellular complexity reflected in an array of different renal functions: from filtering the blood, regulating water homeostasis, production of hormones, to excretion of waste products. These diverse functions are driven by distinct anatomical structures called nephrons. Each nephron comprises more than tens of highly specialized cell types, including abundant epithelial cells supported by vascular, stromal and immune cells^1^. Notably, the function and nomenclature of cells that assemble the nephron depend on their location relative to the main tubular structures: the proximal tubule, loop of Henle, distal convoluted tubules, and the collecting duct^2^.

More than 150 litres of filtrate are reabsorbed by the nephrons in a day. Most of this reabsorption occurs in the proximal tubules, which are primarily located in the cortex, the outermost portion of the kidney. Sodium gradient, generated by the activity of numerous Na+/K+-ATPase channels, drives the transport of salts, water, glucose and amino acids, back to the bloodstream. This process requires large amounts of energy, supplied by the abundant mitochondria. Proximal tubules are thus the most metabolically active structures in the nephron^3–5^. The filtrate then enters the loop of Henle that connects the proximal and distal tubule. Here, fluid and solute transport occurs by diffusion, resulting from the high osmolality gradient in the medulla of the kidney. This gradient is accomplished via differential permeability to water between the descending and ascending portion of the loop^6^. Finally, the filtrate travels through the distal convoluted tubule and collecting duct system, where water is reabsorbed and urine is concentrated^7^ (**Fig. 1A**).

**Figure 1.**
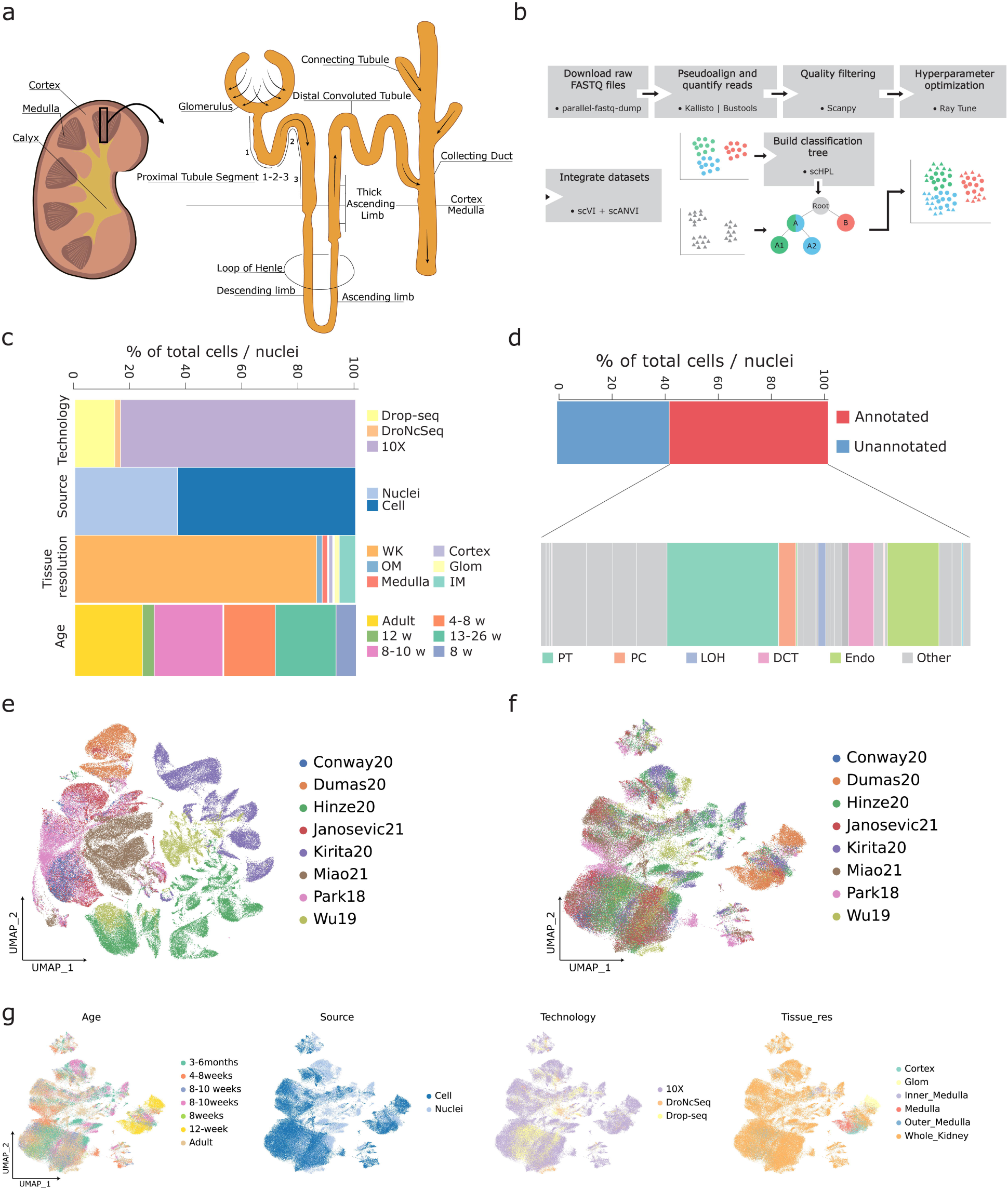
Generation of the mouse kidney atlas from eight independent studies. **a** Schematic of a kidney and a nephron. Arrows indicate the flux of glomerular filtrate through the tubular segments. **b** Workflow guiding the generation of the mouse kidney atlas. Colours represent different hypothetical cell type annotations from two independent studies (A, B and A1, A2) whereas shapes depict originally annotated (circle) or unannotated (triangles) cells and/or nuclei. **c** Metadata information across all datasets. Age of the animals is represented in weeks or months (w, m) when available, otherwise an overall age estimator is provided (Adult). Tissue resolution varied from Whole Kidney (WK), Cortex, Medulla to more selected regions such as Outer Medulla (OM), Inner Medulla (IM) or Glomerulus (Glom). The starting material was either single cell or single nuclei sequenced using Drop-seq, DroNc-Seq or 10x Genomics. **d** Proportion of all annotated cell types across all datasets. Relevant cell types in the nephron are highlighted, namely Proximal Tubule cells (*PT*), Principal Cells (*PC*), Loop of Henle cells (*LOH*), Distal Convoluted Tubule cells (*DCT*) and Endothelial cells (*Endo*). **e** Uniform Manifold Approximation and Projection (UMAP) embedding of all used datasets prior integration. Colours correspond to the different datasets. **f** UMAP visualization of merged datasets following integration and batch correction (see Methods). sCell: single-cell, sNuc: single-nuclei. **g** UMAP representations of the 140K cells and nuclei after integration. Relevant metadata was extracted for each of the datasets. Age of the animals is represented in weeks or months when available, otherwise an overall age estimator is provided (Adult).

Although outstanding efforts to characterize the transcriptomic profiles of the different cell types present in the kidney have been supported by the recent advances in single-cell technologies^8–15^, the identification of markers for distinct and, in particular, rare cell types remains elusive. A substantial portion of the transcriptomic data comes from the proximal tubule and loop of Henle cells, which are one of the largest structures of the nephron^16^. Rare cell populations usually remain undetected and the costs to profile these cells often precludes the studies of less abundant cell populations^17,18^. The aforementioned challenges could be addressed by creating a reference atlas of the kidney, that leverages the vast collection of available cells and nuclei profiled, which can then be used to annotate specific kidney cell types in a supervised manner. Integrating different datasets into a common space can overcome batch effects which occur due to the differences in library preparation protocols and data processing steps. However, in most studies to date, the annotation of cell populations is performed in an unsupervised manner, a process that is time-consuming and involves several refinement iterations^19^. The subjective nature of this approach limits the ability to compare populations across studies due to annotation inconsistencies. Ideally, the reference atlas would account for the different resolutions at which cell populations have been annotated and include a harmonized annotation that allows for a better characterization of rare cell populations.

Here we create an atlas of the adult healthy mouse kidney (MKA: Mouse Kidney Atlas) by integrating and harmonizing annotations from publicly available single-cell and single-nuclei transcriptomic studies. We integrated ∼140.000 cells and nuclei from 59 samples sequenced in eight different studies^8–14^ to generate an atlas that reflects the biological component of the different samples, while accounting for technical differences. We built a hierarchical model of the cell populations present in the healthy mouse kidney, that accurately predicts cell annotations in unlabelled datasets. In addition, MKA allows further integration of new datasets as they become available by relying on a progressive learning approach^20^ **(Fig. 1B**). We show that the new atlas allows the identification of novel and robust markers for both known cell types and previously unexplored rare cell populations.

## Materials and Methods

### Collecting raw data and quantification of reads

All raw fastq files were downloaded from the Sequence Read Archive (SRA) using *parallel-fastq-dump* (v0.6.7; https://github.com/rvalieris/parallel-fastq-dump). Accession numbers and other relevant metadata are provided in **Table 1**.

single-cell and single-nuclei droplet-based sequencing data were aligned and quantified using kallisto/bustools^21,22^ (kb-python v0.26.4) *ref* and *count* wrappers, specifying --workflow nucleus in the case of single-nuclei sequencing experiments. Reads were pseudo aligned to the mouse reference genome GRCm38 downloaded from Ensembl^23^.

### Pre-processing of sequencing data and normalization

Filtered count matrices from Kallisto/bustools were used when the cell count was within a 10k margin from the matrices deposited by the authors. Otherwise, the unfiltered count matrices were loaded, and barcodes were matched between the author’s and the unfiltered set of cells. Count matrices were pre-processed using Scanpy^24^ (v1.8.1). We applied quality filters to all samples, specifically, we only kept cells with a number of detected genes between 200 and 3500. Furthermore, we filtered out cells with more than 50% of counts derived from mitochondrial genes. Samples were merged and normalized for plotting purposes with Scanpy’s normalize_total.

### Integration

The integration was performed using scVI and later refined with scANVI^27,28^ and cell type information. First, as different hyperparameter combinations and model configurations can affect the performance of these models, Ray tune^25^ was used to optimize scVI’s model. Raw counts and batch information were used to test 1000 different hyperparameter combinations. Our search space consisted of model configurations such as continuous and categorical covariates; model hyperparameters such as dropout rate, number of layers and number of latent dimensions; learning hyperparameters such as learning rate and preprocessing steps such as highly variable genes (HVG) filtering and number of hvgs. The objective function to optimize was the silhouette score of both batch and cell type information as implemented in scib^26^. Detailed information and the scripts used to perform these analyses are available at https://github.com/nrclaudio/MKA.

The hyperparameter configuration with the highest silhouette score was then used to train scVI. In our case, we reduced the feature space of our atlas to the top 3000 HVGs. Variable genes were obtained using Scanpy’s highly_variable_genes with the flavour set to seurat_v3. We included the percentage of mitochondrial reads as a continuous covariate in the model and the source of the material (cells or nuclei) as a categorical covariate. The model was initialized with 2 hidden layers, 26 latent dimensions, a dropout rate of 0.096 and the gene likelihood set to a Negative Binomial distribution. The model was then trained for a total of 111 epochs with a learning rate of 0.0013. The obtained model was then used as input for scANVI in order to further improve the latent space representation. We included available cell type annotations and set the unlabeled_category to the set of cells with missing annotations. The scANVI model was trained to a maximum of 20 epochs and with 100 samples per label.

### Dimensionality reduction

After integration and batch-correction, 26 latent dimensions were obtained from the model. These were used as input for the Nearest Neighbor graph calculation using Scanpy’s neighbors function. We further reduced the dimensionality to visualize the data in a 2D UMAP using 26 latent dimensions.

### Similarity metrics

To assess cell population similarity across studies, pairwise similarity measures were computed using sklearn^29^ (v0.23.2) pairwise_distances with the correlation metric. The similarity between two cell populations is reported as 1 – correlation distance between their average normalized transcriptomic profile.

### Integration metrics

Batch and biological conservation metrics were computed using scib (v1.0; https://github.com/theislab/scib). K-nearest neighbour Batch Effect Test^30^ (kBET) was implemented as in Pegasus (v1.4.4; https://github.com/klarman-cell-observatory/pegasus). Batch conservation metrics include graph Local Inverse Simpson’s Index^31^ (LISI), and kBET. In short, these metrics quantify the alignment between the different batch labels in the data. Specifically, kBET examines to what extent the different batches are mixed when neighbourhoods are randomly sampled and LISI captures the diversity of batches within a local neighbourhood of cells.

### Cell type learning and classification

All 26 latent dimensions from the annotated datasets (Park18, Wu19, Miao21, Kirita20, Dumas20 and Janosevic21) along with their original or curated cell type labels were used as input for single-cell Hierarchical Progressive Learning^20^ (scHPL, v1.0.0). For both the original and the curated labels, the classification tree was learnt using a kNN and default values. To classify the cells that were missing annotations, the learnt tree and the latent dimensions from Hinze20 and Conway20 were used as input for scHPL’s predict_labels function.

### Evaluation of the classifier

Using Miao21 data as a validation set, the performance of scHPL trained with Park18, Wu19, Kirita20, Dumas20 and Janosevic21 was evaluated. The classification tree was learnt as described in the previous section. Miao21 labels were predicted and compared to the original curated labels. To measure the accuracy of the prediction, the F1 score (harmonic mean of the precision and recall) was computed using scikit-learn v1.0.1 f1_score function with the average set to micro. This was done for every cell population. Overall F1 scores were computed as the median of F1 scores across populations.

To compare the performance of scHPL trained in our reference with other methods and references, we submitted Miao21 raw count data as a query to Azimuth with the Human Kidney Reference atlas ^32,33^. We kept the quality control filters we applied in our own pre-processing. The *l2*.*annotation* labels were transferred to the query using Azimuth. We also tested the performance of Azimuth’s workflow (In Seurat^34^: Seurat::SCTransform, Seurat::FindTransferAnchors and Seurat::MapQuery) to transfer the labels from our reference to Miao21 query dataset. As described previously, the annotations predicted by Azimuth with the Human Kidney Reference and our own reference were compared to the original curated labels of the query. The accuracy of this prediction was computed as an overall F1 score. For each pair of predicted and original labels, confusion matrices were computed using scHPL’s confusion_matrix function. The row vectors of these matrices were normalized to sum up to 1.

### Differential gene expression analyses and meta-marker discovery

Cell type markers were computed using Scanpy’s rank_gene_groups function with the Wilcoxon rank-sum test. Meta-markers were computed using the MetaMarkers R package (v0.0.1; https://github.com/gillislab/metamarkers). Raw counts were converted to CPM values (as in original work). Markers were computed for each dataset with compute_markers to then obtain meta-markers using make_meta_markers. These two functions are using a Mann-Whitney test per dataset and an aggregation based on meta-analysis of the obtained p-values, respectively. Pareto boundary markers (i.e. markers with high precision and detection rate) were visualized using plot_pareto_markers.

### Comparison with microdissected kidney bulk RNA-seq

TPM values were downloaded from Gene Expression Omnibus (GEO, GSE150338). From a total of 96 cell type bulk RNA-seq libraries, we kept 64 corresponding to the matching cell types in our atlas. TPM values were normalized as log2(TPM + 1). We then visualized the normalized expression of the previously computed cell type markers in the bulk RNA-seq context using pheatmap (https://github.com/raivokolde/pheatmap).

To test the significance of the overlap between the lists of differentially expressed genes (i.e. cell type markers defined by either our kidney atlas or the microdissection study) we used scipy’s v1.5.4 Fisher exact test (fisher_exact) in every cell type present in both the atlas and the microdissection study. We used the list of genes present in the atlas as background in the test. In both cases we considered as significant those genes with an adjusted (using Benjamini-Hochber’s FDR correction) p-value < 0.01. 99% Confidence intervals were computed for the odds ratios obtained in the test. This test evaluates whether a list of significant markers is independent of the list of markers that it is being compared to.

### Data and code availability

Jupyter notebooks and scripts used in the analyses as well as supplemental data are available on Github (https://github.com/nrclaudio/MKA). Interactive visualization and downloading of the kidney mouse atlas are available at cellxgene (https://cellxgene.cziscience.com/e/42bb7f78-cef8-4b0d-9bba-50037d64d8c1.cxg/).

## Results

### Integrated atlas accounts for technical differences among seven independent studies

To create a comprehensive atlas of the healthy mouse kidney we downloaded the raw sequencing data (FASTQ) from eight different studies including a total of 59 samples ^8–14^ (Table 1). To reduce variability in alignment rates between different genetic make-ups, we only included samples with a C57BL/6 background. The raw reads of all samples were processed using the same pipeline and we recovered 140.000 cells and nuclei after filtering low quality cells and nuclei (see Methods section for details). The samples included in this study differ in single-cell technology, source of material, tissue resolutions and age of sacrifice **(Fig. 1C)**. Approximately 40% of the cells and nuclei included in this study were missing computer-readable annotations **(Fig. 1D)**. These differences can be visualized in the Uniform Manifold Approximation and Projection (UMAP) of the data **(Fig. 1E)**, where source-specific populations were identified. To resolve these batch effects, we used scvi-tools’ scVI and scANVI ^27,28^. After integration, the aligned compendium demonstrates that the different data sources have been properly aligned and no metadata is driving the differences observed in the UMAP space **(Figs. 1F and G)**. We used the k-nearest neighbour Batch Effect Test (kBET) and graph Local Inverse Simpson’s index (LISI) metrics to quantify the quality of batch correction. Both kBET and LISI increased from a value of zero for the unintegrated data to a kBET of 0.024 and a LISI of 0.27, indicating proper alignment of batches.

### Integration highlights annotation inconsistencies across studies

After integration, we investigated cell population annotations across the four data sets for which annotations were available or were manually annotated (Park18, Wu19, Kirita20, Miao21, Dumas20 and Janosevic21). The six datasets varied significantly in the resolution and the ontology used to annotate distinct cell populations. Only two cell populations were common between the five studies (Dumas20 only surveyed endothelial cells) based on the set of author’s annotated terms, with most of the annotations being dataset-specific **(Fig. 2A)**. For example, collecting duct intercalated cells (*IC*) can be further classified into Type A (*ICA*) or Type B (*ICB*), depending on the expression and localization of Slc4a1 in the membrane and the presence of a transport protein called pendrin, encoded by the *Slc26a4* gene. Whereas *ICA* cells lack pendrin and acidify the urine by excreting H^+^, *ICB* cells have pendrin and secrete OH^-^ equivalents^35^. Another example are Proximal Tubule Cells (*PT*). While certain studies identify *PT* cells, some others further classify these cells in three different segments (*PTS1, PTS2* or *PTS3*) depending on their location along the nephron **(Fig. 1A and 2B)**. These differences in ontology, together with the distinct annotation resolutions, highlight the subjective nature of unsupervised cell type annotation and the need for an integrated and comprehensive view of cell heterogeneity in the kidney.

**Figure 2.**
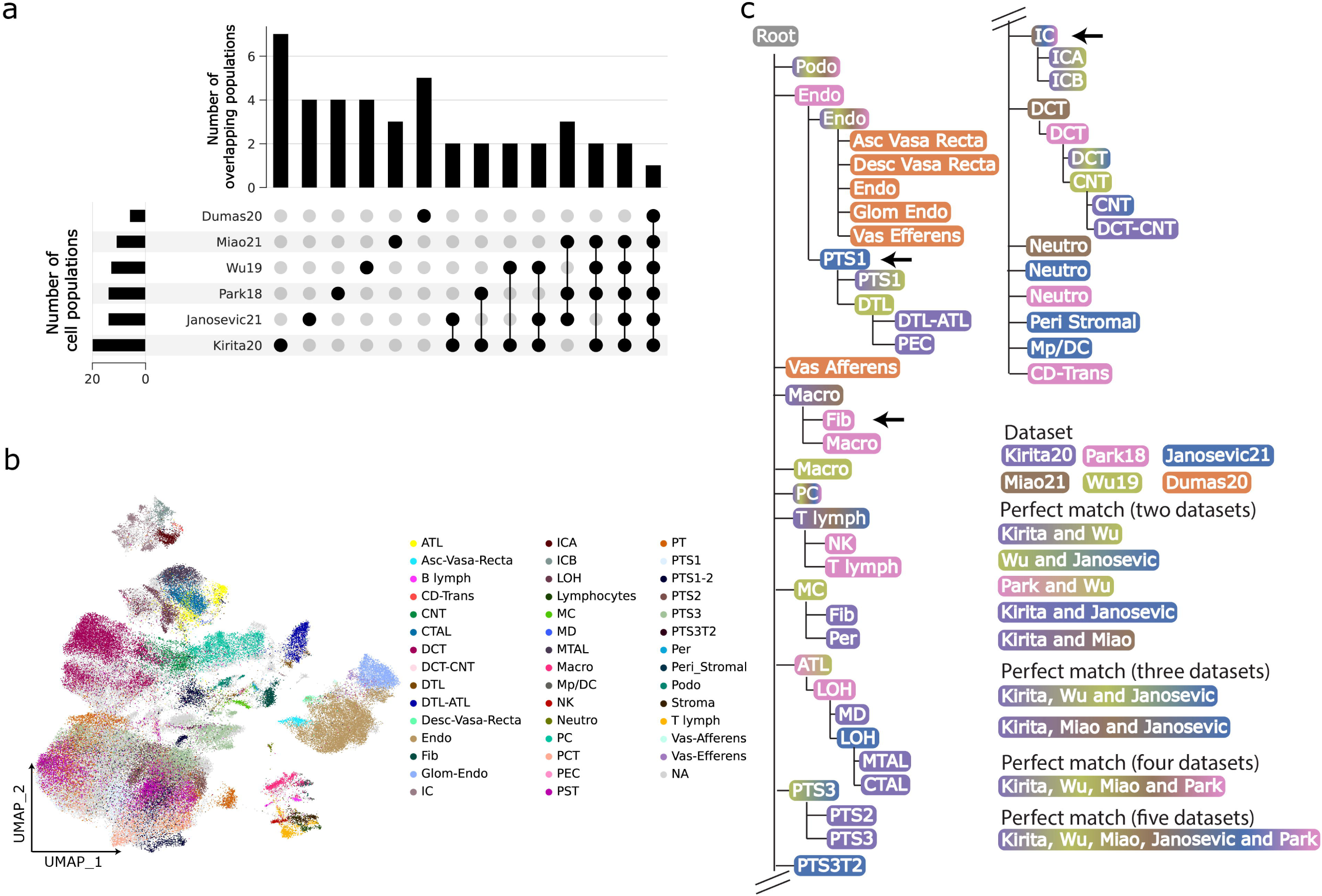
Learned classification tree from independently annotated datasets. **a** UpSet plot visualizing the intersection and the number of common cell type annotations between the different datasets. Disconnected dots correspond to the number of unique cell types in each dataset, while connected dots represent the intersection between the datasets. Additionally, the number of cell types identified in each study is plotted alongside each dataset. **b** UMAP representation of the originally annotated cell types across all datasets. **c** Learned classification tree applying a k-Nearest Neighbor (kNN) classifier on the six annotated datasets. The colour(s) of the tree nodes correspond to the supporting dataset(s). Arrows mark both inaccurate placements in the tree (*Fib, PTS1*) and cell types that can be further annotated to increase resolution (*PT*: *PTS1, PTS2, PTS3*; *IC*: *ICA, ICB*). *IC*: Intercalated Cell, *ICA*: Intercalated Cell Type A, *ICB*: Intercalated Cell Type B, *Endo*: Endothelial Cell, *Fib*: Fibroblast, *Macro*: Macrophage, *B lymph*: B lymphocyte, *Stroma*: Stroma cell, *NK*: Natural Killer, *T lymph*: T lymphocyte, *PT*: Proximal Tubule, *PTS1*: Proximal Tubule Segment 1, *PTS2*: Proximal Tubule Segment 2, *PTS3*: Proximal Tubule Segment 3, *PTS3T2*: Proximal Tubule Segment 3 Type 2, *PC*: Principal Cell, *PEC*: Parietal Epithelial Cell, *Per*: Pericyte, *DCT*: Distal Convoluted Tubule, *ATL*: Ascending Thin Limb of Henle, *MD*: Macula Densa, *LOH*: Loop of Henle, *CTAL*: Thick Ascending Limb of Henle in Cortex, *MTAL*: Thick Ascending Limb of Henle in Medulla, *CNT*: Connecting Tubule, *Podo*: Podocyte, *DTL*: Descending Thin Limb of Henle, *MC*: Mesangial Cell, *Neutro*: Neutrophil, *Asc-Vas-Recta*: Ascending Vasa Recta, *Desc-Vas-Recta*: Descending Vasa Recta, *Glom Endo*: Glomeruli Endothelial.

To overcome these challenges, we used single-cell Hierarchical Progressive Learning (scHPL)^20^, a method that automatically infers cell hierarchies from annotated datasets and builds a classification tree that can be used to classify unlabelled cells. We used scHPL to build a cell hierarchy and capture the relationships between kidney cell populations from the six annotated datasets. Perfect matches were found between cell populations across five (Principal Cells, *PC*), four (*Endothelial* and *Podocytes*) three (Distal Convoluted Tubule, *DCT* and *T lymphocytes*) and two datasets (*PTS1, PTS3*, Ascending Thick Limb of Henle, *ATL, Macrophages, ICA, ICB* and *CNT*). Not a single cell population matched across all six datasets **(Fig. 2C)**. On the other hand, some cell populations are misplaced in the tree. For example, the *fibroblasts* from Park18 (hereafter *cell type*_dataset_) are placed under *Macrophages*_Kirita20, Miao21_ another example are *PTS1*_Janosevic21_ cells which are placed under *Endothelial*_Kirita20, Wu19, Park18, Miao21_ cells. Other populations are lacking available resolution. For example, *IC*_Park18, Miao21_ cells, which appear as parent node of *ICA*_Kirita20, Wu19_ and *ICB*_Kirita20, Wu19_ cells. *Peri Stromal*_Janosevic21_ cells Moreover, some cell populations are missing in the final tree because they have been rejected (e.g., *PT*_Park18_ cells and *PCT*_Miao21_ and *PST*_Miao21_ cells could not be classified).

### Manual curation of annotations significantly improves hierarchy learning

To refine the cell tree constructed by scHPL, we performed a manual curation of the original cell population annotations **(Figs. S1 to S3)**. The initial tree constructed by scHPL indicates that *Stroma*_Miao21_ cells have similar transcriptomic profiles to *T lymphocytes*_Park18_ **(Fig. 2C)**, which is supported by their overlap in the UMAP and the high similarity of their average expression profile **(Figs. S1A to S1C)**. This observation was supported by the expression of T lymphocyte markers^36,37^ (*Cd4, Cd8a, Cd28*) in cells annotated as *Stroma* **(Fig. S1D)**. In addition, we compared the expression of *Cd4, Cd8a* and *Cd28* in *Stroma*_Miao21_ cells, *T lymphocytes*_Park18_ and Kirita20 non-immune cell types **(Fig. S1E)**. As expected, *Stroma*_Miao21_ cells share the expression of these markers with *T lymphocytes*_Park18_ but not with non-immune populations. A similar scenario applies to *Fibroblasts*_Park18_, which are placed under the Macrophages node **(Figs. 2C and S1E)**. We checked whether these cells might have been mislabelled by visualizing the expression of M1-M2 Macrophage markers^36,38^ (*Cd68, H2-Ab1* and *Il4r*) **(Fig. S1F and S1G)**. Based on these observations, we re-annotated *Stroma*_Miao21_ and *Fibroblasts*_Park18_ to *T lymphocytes* and *Macrophages*, respectively.

We then evaluated the location of *PT* cells in the tree, which can be further classified in different segments (Segments 1, 2, 3 and 3 type 2; *PTS1, PTS2, PTS3, PTS3T2*). These segments, where numerous transporters regulate endocrine functions, can be morphologically identified close to the nephron’s glomerulus ^5^. Park18 included the lower resolution term *PT* **(Fig. S2A)** and Miao21 annotations included the terms Proximal Straight Tubule (*PST*) and Proximal Convoluted Tubule (*PCT*) **(Fig. S2B)**, Wu19 grouped *PTS1* and *PTS2* cells together **(Fig. S2C)** and Kirita20 did not include *PTS3T2* **(Fig. S2D)**. To re-annotate these cells as *PTS1, PTS2, PTS3* or *PTS3T2*, we used unsupervised clustering and visualized known markers to rename the resulting cell populations. The visualized markers were *Slc5a12, Cyb5a, Slc27a2* and *Cyp7b1* for *PTS1, PTS2, PTS3* and *PTS3T2*, respectively^15,39^. In the case of *PTS1-2*_Wu19_, the population was matched to *PTS1*_Kirita20_ during training of scHPL. We re-annotated *PTS1-2*_Wu19_ as *PTS1*_Wu19_.

We refined the annotation of *IC*_Park18_, *IC*_Miao21_ and *Endothelial*_Park18_ cells following the same strategy as described above **(Figs. S3A and S3B)**. *IC*_Park18_ and *IC*_Miao21_ cells were re-annotated as either *ICA* or *ICB* based on the expression of *Slc4a* (ICA marker) and *Insrr* (ICB marker) in the unsupervised clusters **(Fig. S3C)**. *Endothelial*_Park18_ cells were originally re-annotated as Descending Thin Limb of Henle (*DTL*) by the original authors in the manuscript, but this correction was missing from the annotations provided with the dataset. Therefore, we similarly refined the annotation based on the expression of *Slc14a2* (DTL marker) and *Adgrl4* (Endothelial marker) **(Fig. S3D)**.

Following these annotation refinements, we applied scHPL to rebuild the cell tree and re-trained the classifier on the new tree. The new tree correctly captures the expected cell hierarchy, placing similar populations within the same node **(Fig. 3A)**. This is exemplified by the identification of the Medullary and Cortically Thick Ascending Limb of Henle (*MTAL* and *CTAL*) as child nodes of Loop of Henle (LOH) cells **(Fig. 1A)**. This shows the ability of the hierarchical model to group functionally and morphologically related cell types in the nephron. In this case, perfect matches between datasets were more easily identified, likely due to the lower number of rejected cells while training.

**Figure 3.**
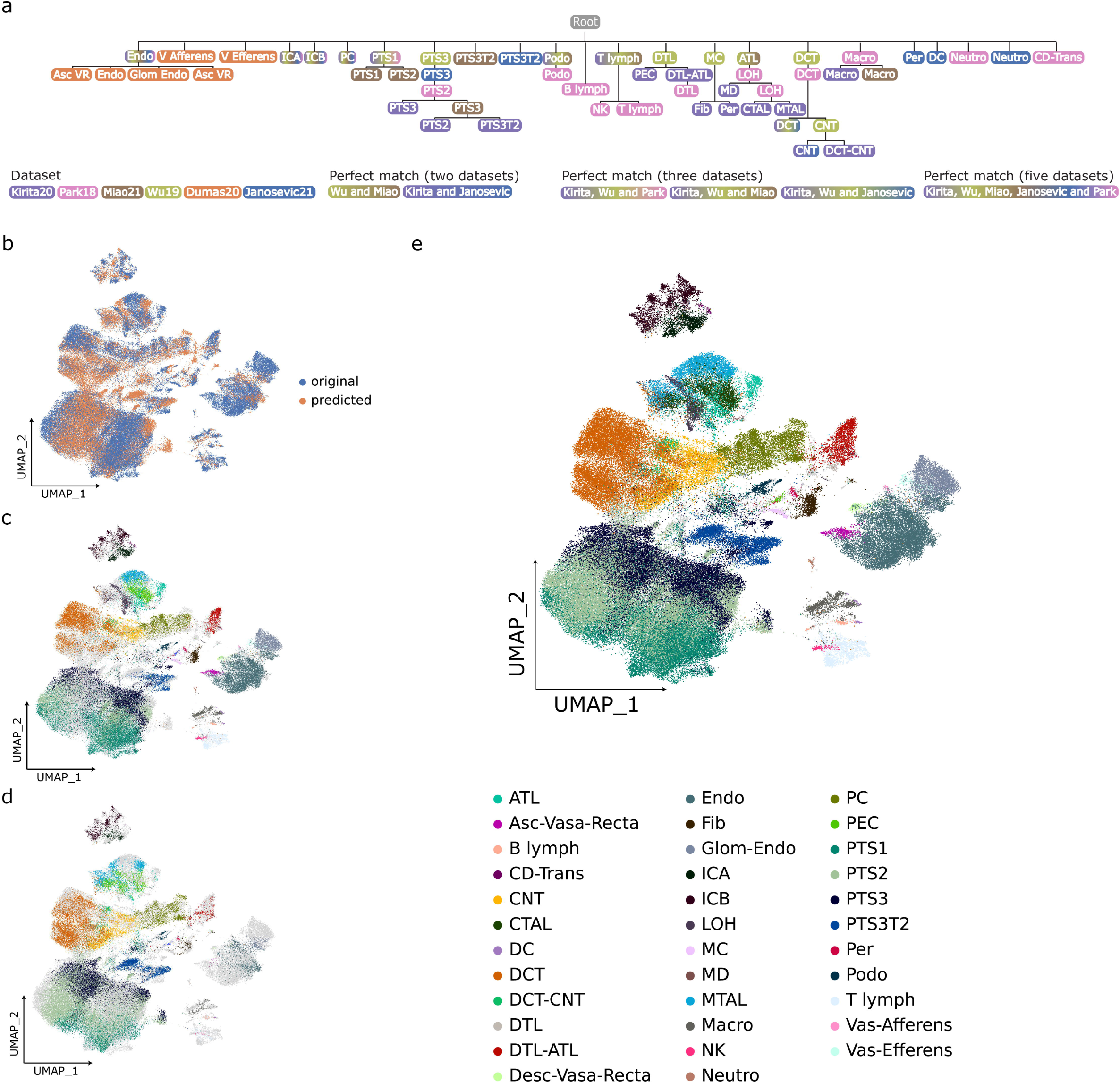
Hierarchically defined kidney mouse atlas. **a** Learned classification tree using a linear SVM on the four annotated datasets after manually curating misclassified cell types in the tree or inaccurate annotations. Tree nodes are coloured by the supporting dataset or datasets in case of two or more cell populations matching. **b** UMAP plot visualizing cells used to train the classification tree (blue) and cells for which cell type was not known (orange). **c** UMAP representation of annotated cell types for all datasets. **d** UMAP embedding of predicted cell types. **e** UMAP plot combining predicted and available annotations resulting in *the integrated mouse kidney atlas. IC*: Intercalated Cell, *ICA*: Intercalated Cell Type A, *ICB*: Intercalated Cell Type B, *CD-Trans*: Collecting Duct Transitional cell, *Endo*: Endothelial Cell, *Fib*: Fibroblast, *Macro*: Macrophage, *B lymph*: B lymphocyte, *Stroma*: Stroma cell, *NK*: Natural Killer, *T lymph*: T lymphocyte, *PT*: Proximal Tubule, *PTS1*: Proximal Tubule Segment 1, *PTS2*: Proximal Tubule Segment 2, *PTS3*: Proximal Tubule Segment 3, *PTS3T2*: Proximal Tubule Segment 3 Type 2, *PC*: Principal Cell, *PEC*: Parietal Epithelial Cell, *Per*: Pericyte, *DCT*: Distal Convoluted Tubule, *ATL*: Ascending Thin Limb of Henle, *MD*: Macula Densa, *LOH*: Loop of Henle, *CTAL*: Thick Ascending Limb of Henle in Cortex, *MTAL*: Thick Ascending Limb of Henle in Medulla, *CNT*: Connecting Tubule, *Podo*: Podocyte, *DTL*: Descending Thin Limb of Henle, *MC*: Mesangial Cell, *Neutro*: Neutrophil, *Asc-Vas-Recta (Asc VR)*: Ascending Vasa Recta, *Desc-Vas-Recta (Desc VR)*: Descending Vasa Recta, *Glom Endo*: Glomeruli Endothelial, *V afferens*: Vas Afferens, *V Efferens*: Vas Efferens.

Next, we used the refined classification tree from scHPL **(Fig. 3A)** to predict the cell type annotations for cells and nuclei from the remaining two unlabelled studies (Conway20 and Hinze20) **(Figs. 3B to 3D)**. After merging the predicted and original labels, we obtained the final fully annotated adult healthy kidney atlas (MKA) **(Fig. 3E)**. The complete overview of the cell population shows that the integration process preserved the shared biological component between the different studies.

Due to the lack of labels for these four studies, we could not perform a quantitative analysis of the obtained labels. To confirm our annotations, we visualized known markers for the major cell types in the nephron. Namely *PT, Podocytes, DTL, ATL, MTAL, CTAL, DCT, CNT, ICA, ICB* and *Endothelial* cells **(Figs. S4A to S4D)**. Moreover, we compared the cellular composition of each study to that reported in the original studies. We found that the proportion of all predicted Proximal Tubule Cells, i.e., *PTS1, PTS2, PTS3* and *PTS3T2*, matches the proportion described in the original publications, 77% in Conway20 and approximately 60% in Hinze20 **(Fig. S4E)**. The same applies to *MTAL, CTAL* and *DCT* in the MKA. These cell populations were described as *LOH/DCT* in Conway20 and *TAL* in Hinze20 with a proportion of approximately 7% and 10% respectively **(Fig. S4E)**.

The MKA allowed us to annotate these datasets at a higher resolution than originally reported. For example, in Conway20 they annotated 15 cell types. We now identify 28 distinct populations, providing further resolution for annotations such as *LOH/DCT* (*MTAL, CTAL* and *DCT* in the MKA) or *CD* (*CD-Trans, ICA, ICB* and *CD* in the MKA). We also identify previously overlooked important cell populations such as *PC* **(Fig. S4B)**. Another example is Hinze20, in which MKA identified 25 subpopulations among the original set of 10 cell types, including: *PTS1, PTS2, PTS3* and *PTS3T2* instead of *PT*; *ICA* and *ICB* instead of *CD-IC*; and *DTL* and *ATL* instead of *TL* (Thin Limbs) **(Fig. S4A)**.

### scHPL and the mouse kidney atlas accurately classifies unseen cells

To evaluate the accuracy of the scHPL classifier, we performed a leave-one-dataset-out experiment. We chose the Miao21 dataset as the test set and trained the scHPL classifier on Park18, Kirita20, Wu18, Dumas20 and Janosevic21 (hereafter called MKA*), with the same parameters as defined earlier in this manuscript. The resulting tree **(Fig. 4A)** was then used to train the scHPL classifier and predict the labels of cells from Miao21. Most of the original annotations from the dataset were accurately predicted by the scHPL classifier with median F1 score of 0.74 **(Figs. 4B and 4F)**. Moreover, scHPL further classified cells at a higher resolution compared to the low-resolution labels present in the original dataset **(Figs. 4E)**. For example, in the original study, Miao21 identified *LOH* cells, which scHPL can classify into *MTAL* and *CTAL*, the two major cell types present in the thick ascending limb of the Loop of Henle. Notably, some cells and nuclei were assigned to the root node (i.e. unclassified). For example, *Neutrophils* were mostly rejected **(Fig. 4B)**. This is not surprising, since there were only 26 *Neutrophil* cells in the training data (i.e. MKA*), which inevitably led to a poor performance in predicting *Neutrophils* in Miao21. On the other hand, one of the most abundant cell types in the training data (*DCT* with 4559 cells and nuclei) is correctly predicted 90% of the time (*DCT* and *DCT-CNT*). scHPL can reject cells due to the lack of cells from a specific population during training, e.g. neutrophils. But rejection can also mean that the query dataset includes novel cell populations not seen during training. In the latter case, rejected cells assigned can be further characterized and annotated to update the cellular knowledge stored in MKA.

**Figure 4.**
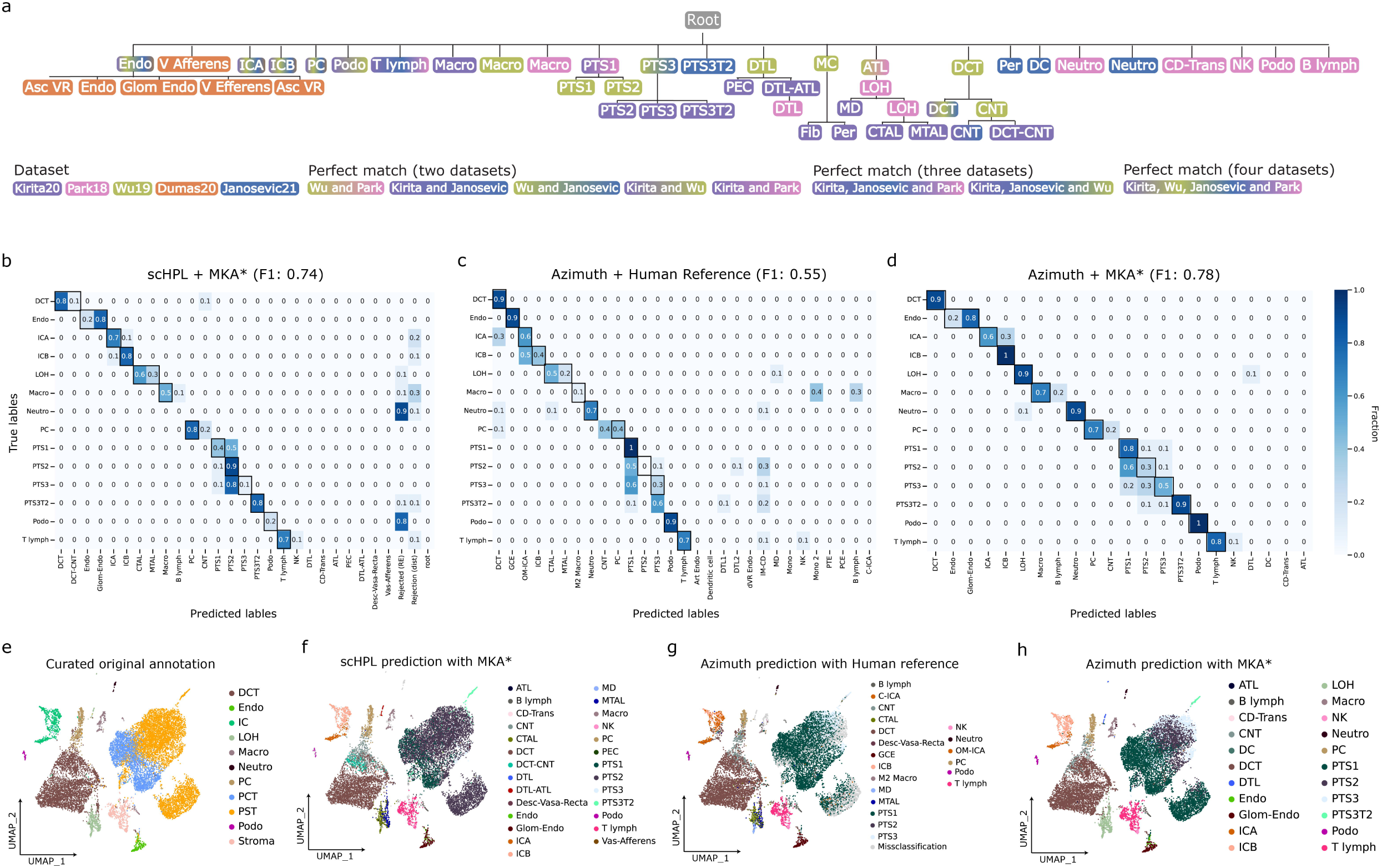
Evaluation of the scHPL classifier. **a** Learned classification tree when using a k-Nearest Neighbor (kNN) classifier on five of the six annotated datasets MKA* (Miao21 dataset was excluded). The colour(s) of the tree nodes represent the agreement with the supporting dataset(s) **b-d** Confusion matrices normalized by class support size, computed using the predicted annotations by scHPL and our Atlas reference **b**, the transferred labels from Azimuth’s human kidney reference^11^ **c** or the transferred labels from Azimuth’s using our reference **d**. Higher values indicate higher agreement between predicted and true cell labels. **e-h** UMAP plot of the Miao21 dataset coloured by the original cell types (after manual re-annotation) **e**, by the predicted cell types from the learned classification tree **f**, by the transferred cell types from the Azimuth human reference **g** and by the transferred labels using Azimuth with our Atlas reference **h**. *IC*: Intercalated Cell, *ICA*: Intercalated Cell Type A, *ICB*: Intercalated Cell Type B, *Endo*: Endothelial Cell, *Fib*: Fibroblast, *Macro*: Macrophage, *B lymph*: B lymphocyte, *Stroma*: Stroma cell, *NK*: Natural Killer, *T lymph*: T lymphocyte, *PT*: Proximal Tubule, *PTS1*: Proximal Tubule Segment 1, *PTS2*: Proximal Tubule Segment 2, *PTS3*: Proximal Tubule Segment 3, *PTS3T2*: Proximal Tubule Segment 3 Type 2, *PC*: Principal Cell, *PEC*: Parietal Epithelial Cell, *Per*: Pericyte, *DCT*: Distal Convoluted Tubule, *ATL*: Ascending Thin Limb of Henle, *MD*: Macula Densa, *LOH*: Loop of Henle, *CTAL*: Thick Ascending Limb of Henle in Cortex, *MTAL*: Thick Ascending Limb of Henle in Medulla, *CNT*: Connecting Tubule, *Podo*: Podocyte, *DTL*: Descending Thin Limb of Henle, *MC*: Mesangial Cell, *Neutro*: Neutrophil, *Asc-Vas-Recta (Asc VR)*: Ascending Vasa Recta, *Desc-Vas-Recta (Desc VR)*: Descending Vasa Recta, *Glom Endo*: Glomeruli Endothelial. *GCE*: Glomerular Capillary Endothelial. *OM-ICA:* Outer Medullary *ICA, C-ICA*: Cortical *ICA, Art Endo*: Afferent / Efferent Arteriole Endothelial, *PCE*: Peritubular Capilary Endothelial cell, *PTE*: Peritubular Endothelial cell.

Next, we sought to compare the performance of scHPL with Azimuth^32^, a widely used pipeline to automatically annotate cells based on Seurat. At the moment, Azimuth only includes a human kidney reference atlas^33^. We performed two experiments in which we used Azimuth to predict the labels of our query dataset (Miao21). First, we evaluated the performance of Azimuth using the provided human reference in predicting mouse cells from Miao21. Despite having a wider array of cell population (46 populations), Azimuth misclassified many cells with a median F1 score of 0.55 **(Figs. 4C and 4G)**. For instance, *PC* cells were classified as *CNT* or *DCT* 50% of the time. This is a high rate of misclassification considering that these are two very distinct cell types specialized in different functions in the nephron. These misclassifications can be due to the lack of a rejection option in Azimuth or differences in the cell type-specific transcriptomic profiles between human and mouse kidney, or a combination of both factors. Second, we tested Azimuth’s performance using our partial reference atlas (MKA*), which was constructed from Park18, Kirita20, Wu19, Dumas20 and Janosevic21. Using Azimuth with MKA* as a reference, we were able to accurately classify cells in the query data with a median F1 score of 0.78 **(Figs. 4D and 4H)**. This result indicates that the low performance of Azimuth compared to scHPL is mainly due to the use of a human reference to classify mouse cells. In order to understand how different populations contributed to the F1 score, we computed a F1 score per cell type and model **(Fig. S5)**. Three of the fourteen populations included were accurately classified (F1 > 0.8) across the different validation experiments. In the case of the MKA*+scHPL experiment, the number of accurately classified populations increases to eight out of fourteen. Despite Proximal Tubule cells (*PTS1, PTS2, PTS3* and *PTS3T2*) being the most abundant cell type in the nephrons^36^, we saw a lot of variation in the classification accuracy of these populations. In the training data (MKA*), PT cells account for 45% of the total number of cells and nuclei. *PTS3T2* cells (3%) are accurately predicted when using MKA* and either scHPL or Azimuth. This can be explained by the lack of this population in the human kidney reference available. *PTS1* and *PTS2* cells (10% and 17%) display a high degree of F1 score variability across the different experiments **(Fig. S5)**. This is expected, as segments 1 and 2 can be identified morphologically but have almost identical functionality in the nephron^40^. As a consequence, their transcriptomic profiles are highly overlapping, which has led to several authors considering them a single cell type^11,14^. None of the reference and classifier combinations we tested accurately classifies both segments. *PTS3* is the least abundant cell type in the nephron *PTS3* cells have the highest accuracy score when using the MKA* with Azimuth. Even in this case, 50% of the time *PTS3* cells are misclassified as either *PTS1* or *PTS2* **(Fig. 4D)**.

### Mouse Kidney Atlas facilitates the identification of robust cell population markers

Technological limitations in single-cell transcriptomics result in a high proportion of unmeasured genes leading to low replicability of cell type markers across different studies. We capitalized on the large collection of cells and nuclei from diverse samples in MKA to identify replicable cell population markers.

Based on MKA, we identified *meta-markers*, which are genes that have a high detection rate and are reliable markers for a given cell population across different datasets (see Methods for details). The resulting set of meta-markers per cell type included previously known markers (e.g. *Slc12a1* for both *MTAL* and *CTAL*, and *Slc12a3* for *DCT*) as well as novel candidates (e.g. *Bst1* for *DTL* or *Rhcg* for *CNT*) **(Fig. 5A and Supplementary Data 1)**.

**Figure 5.**
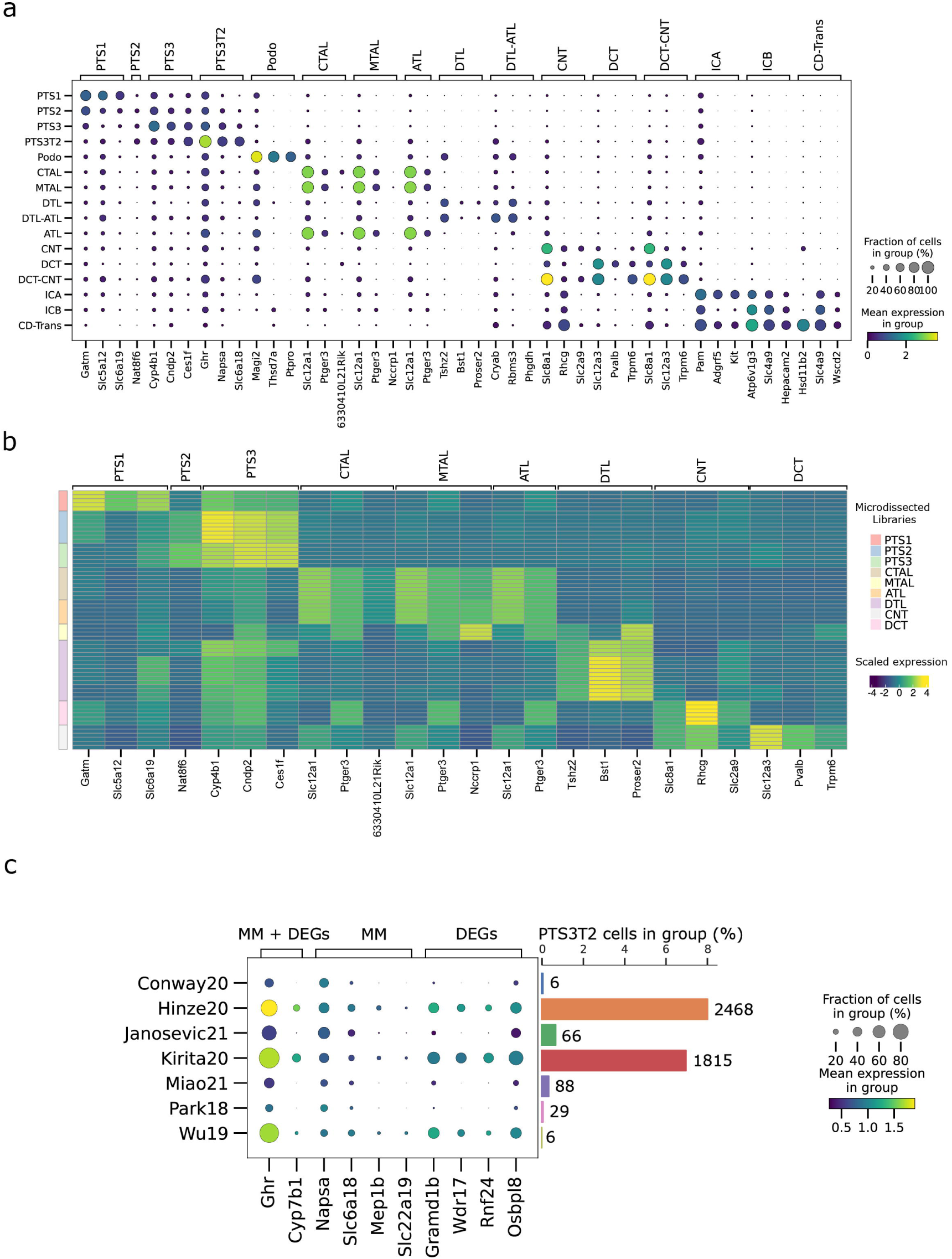
Joint downstream analyses highlight known cell type markers and help define meta-markers across studies. **a** Dotplot of the top 3 meta-markers (when their recurrence is equal or greater than 2) per cell type across datasets. Values sorted by fold change of detection rate and auroc. **b** Heatmap showing the scaled and normalized transcript per million (TPM) expression of the top meta-markers in the micro dissected kidney bulk-RNAseq libraries^39^. Only matching cell types between the two experiments were kept. Columns represent the RNA-seq libraries, rows correspond to genes. Both rows and columns are annotated by cell type. In the case of rows, the annotation corresponds to the cell type from which these markers were identified in the atlas. For the columns, the annotations are the different regions from which the RNA-seq libraries are derived from. **c** Dotplot of the top 5 meta-markers (when their recurrence is equal or greater than 2; sorted by fold change of detection rate and auroc) and the top 5 DEGs for *PTS3T2*. Barplots show the number of *PTS3T2* cells in each dataset both in relative (% of total cells in the dataset) and absolute terms (total number of cells on top of each bar). *IC*: Intercalated Cell, *ICA*: Intercalated Cell Type A, *ICB*: Intercalated Cell Type B, *Endo*: Endothelial Cell, *Fib*: Fibroblast, *Macro*: Macrophage, *B lymph*: B lymphocyte, *Stroma*: Stroma cell, *NK*: Natural Killer, *T lymph*: T lymphocyte, *PT*: Proximal Tubule, *PTS1*: Proximal Tubule Segment 1, *PTS2*: Proximal Tubule Segment 2, *PTS3*: Proximal Tubule Segment 3, *PTS3T2*: Proximal Tubule Segment 3 Type 2, *PC*: Principal Cell, *PEC*: Parietal Epithelial Cell, *Per*: Pericyte, *DCT*: Distal Convoluted Tubule, *ATL*: Ascending Thin Limb of Henle, *MD*: Macula Densa, *LOH*: Loop of Henle, *CTAL*: Thick Ascending Limb of Henle in Cortex, *MTAL*: Thick Ascending Limb of Henle in Medulla, *CNT*: Connecting Tubule, *Podo*: Podocyte, *DTL*: Descending Thin Limb of Henle, *MC*: Mesangial Cell, *Neutro*: Neutrophil, *Asc-Vas-Recta (Asc VR)*: Ascending Vasa Recta, *Desc-Vas-Recta (Desc VR)*: Descending Vasa Recta, *Glom Endo*: Glomeruli Endothelial.

In a recent study, Chen and colleagues^39^ surveyed the gene expression of microdissected kidney segments using 64 bulk RNA-seq samples. These segments are identified morphologically and ideally contain a single cell population each. To validate our newly identified meta-markers, we confirmed the expression of the top three meta-markers that appeared in at least two per cell population in the bulk RNA-seq samples. These correspond to *PTS1, PTS2, PTS3, CTAL, MTAL, ATL, CNT, DCT* and *DTL* **(Fig. 5B)**. Furthermore, we found a significant overlap between the MKA-based meta-markers and the microdissection-defined markers **(Table 2)**.

To highlight the value of MKA and the meta-markers we identified, we investigated rare, understudied cell populations. First, we characterized a recently-described cell type, *PTS3T2* cells^15,41^. Together with *PTS3, PTS3T2* cells are thought to play an important role in the kidney injury process^42^. However, the few available marker genes for *PTS3T2* are based on unsupervised clustering of scRNA-seq studies^15^ and are yet to be validated. Within our MKA-based meta-markers for *PTS3T2*, we identified previously known markers, such as *Slc22a13*, as well as novel markers: *Ghr* or *Mep1b* **(Fig. S6A)**. *Ghr* has been previously associated with chronic kidney disease^43^, whereas *Mep1b* plays a role in acute kidney injury, with *Mep1b*^-/-^ mice showing improved renal function compared to WT mice^44^. We compared the expression of the top five *PTS3T2* meta-markers with the top five *PTS3T2* differentially expressed genes in the MKA **(Fig. 5C and Supplementary Data 2)**. Meta-markers such as *Slc6a18* and *Napsa* displayed a robust expression pattern across the non-endothelial datasets (excluding Dumas20). However, DEGs such as *Gramd1b, Wdr17, Rnf24* and *Osbpl8* were expressed mostly at datasets with the highest number of *PTS3T2* cells, lacking replicability across studies. This was the case for the meta-markers *Mep1b* and *Slc22a19* too. *Ghr*, which encodes the growth hormone receptor, was identified as both a meta-marker and a DEG with detectable expression in all datasets. However, *Ghr* is a significant DEG in 30 of the 36 cell populations included in the MKA **(Supplementary Data 1)**, indicating that *Ghr* expression is not specific. *Cyp7b1* is also identified as both a meta-marker and DEG but its expression pattern is biased towards Hinze20, Kirita20 and Wu19.

Although single-cell studies usually aim to describe discrete cell types, kidney’s nephrons are tubular structures formed by a continuum of epithelial cells. Due to this, cells with mixed transcriptomic profiles are likely to be sequenced^9^. We set out to define meta-markers that are known for the cell types that are part of the mixed population, but also to identify novel markers of transitional cell types. In the case of *DCT-CNT* **(Fig. S6B)**, meta-markers included known markers for both *DCT*^39^ (*Slc12a3* and *Slc8a1*) and *CNT*^39^ (*Trpm6*) cells. Novel markers for this mixed population included *Acss3* and *Ltc4s*.

*Cryab*, a known marker for *ATL* cells, is identified as a meta-marker for *ATL-DTL* cells **(Fig. S6C)**. Other not previously known meta-markers include *Rbms3, Phgdh* and *Slc4a11*. Some of these genes have already been implicated in kidney biology. For instance, *Phgdh* has been identified as a treatment target in kidney cell carcinoma in patients resistant to HIF2α antagonists^45^. *Slc4a11* is known to be expressed in *DTL* cells, although expression has been described only in the medullary part of the kidney^46^.

Next, we investigated the novel collecting duct transitional cell population (*CD-Trans*), which was described by Park and colleagues^14^. *CD-Trans* cells have a distinct transcriptomic profile from *ICA* and *ICB* cells which play an important role in the regulation of acid-base homeostasis^47^. Further understanding of *CD-Trans* cells have been hampered by their low abundance in the kidney, often being masked by other cell types, such as Proximal Tubule cells. In MKA, CD-Trans cells were identified in four datasets (Park18, Miao21, Janosevic21 and Conway20) after annotation of the full atlas with 60 cells in total. Our meta-marker list for *CD-Trans* cells includes *Hsd11b2, Slc4a9, Wscd2* and *B3gnt7* which were found to be highly accurate and able to confidently classify cells as *CD-Trans* **(Fig. S6D)**. Kidney-specific *Hsd11b2*^-/-^ mice show systemic salt-dependent hypertension^48^.

## Discussion

The maturity of single-cell and single-nuclei transcriptomics becomes apparent by the ever-increasing number of publications applying these technologies^49,50^. Although this has given rise to a vast collection of publicly available cellular transcriptomes, researchers continue to analyse their work in an isolated environment, often without considering the data from other reports. As it has been recently noted in the literature^40^, the relationships between the populations defined in kidney single-cell studies are not clear and integrative studies are needed. Here, we integrate cells and nuclei from eight independent studies **(Table 1)** to create the first mouse kidney atlas. We demonstrate that, despite between-sample biological and technical differences, our atlas establishes a robust and comprehensive view of the cell heterogeneity present in the mouse kidney.

A major challenge in single-cell analyses is cell type annotation. Usually, cell types are annotated based on the expression of marker genes in unsupervised clusters. Clustering algorithms require the tuning of hyperparameters, leading to a subjective choice on the number of clusters. This is aggravated by the possible presence of new (sub)cell types in the dataset, which usually causes over-clustering ^51^. This introduces subjectivity to the analysis, ultimately leading to incomplete and ambiguous annotations between studies. We highlight these inconsistencies in the case of the mouse kidney using a scHPL, supervised hierarchical machine learning model. By refinement of these annotations and further cell type learning, we improve the atlas reference transcriptome, accurately capturing consensus cell identities across studies. An important feature of such a model is its ability to capture the different resolutions at which cell types have been annotated. For example, some studies limit their labelling to *LOH* cells while others further classify these cells as *MTAL* or *CTAL*^9,14^. In our work we convey a hierarchically defined atlas, further characterizing the variety of cell types present in the healthy mice kidney **(Fig. 3E)**. In consequence, we identify 35 distinct cell types, including both high- and low-resolution annotations. Unfortunately, our atlas cannot predict, with full accuracy, all cell types in the kidney. This limitation is not exclusive to this organ, as supervised cell classification remains a challenge for all tissues. It is often due to the lack of a precise definition of cell types, lack of robust markers, technical limitations, and sampling variability^52^. In addition, renal plasticity and the ability of renal cells to switch cell type might generate some less defined cells^53,54^. In our work we highlight the common misclassification of *PTS1, PTS2* and *PTS3* cells by different methods **(Figs. 4B to 4D)**. Although functional differences between the segments are known, the different segments have traditionally been identified based on cell ultrastructure^55^. This results in their transcriptomes being too similar, rendering these cells hard to classify computationally. We would like to note, however, that by extensive integration of datasets we can largely overcome these shortcomings, as we have demonstrated in the present work. As the field develops, and clearer definitions are proposed, the inclusion of more datasets into our atlas will further enhance cell type identities and classification.

The importance of our work is further highlighted by the pressing need to develop novel therapies for kidney failure. Kidneys are the most frequently transplanted organ. Due to the increasing prevalence of chronic kidney diseases in the population, demand exceeds the number of available donors. and strategies based on renal (stem) cells are being investigated. On grounds of these and other shortcomings, as noted in the literature^56^, an understanding of the cell heterogeneity present in the kidney is needed in order to develop much-needed therapies. The efficacy of these will depend on the cell type-specific expression and activity of pathways ^57^. Despite this, the knowledge of cell types, its markers and the molecular mechanisms and pathology underlying these diseases at the single-cell level is still incomplete. For example, a recent study shows that *CNT* cells can display a partial *DCT* phenotype^58^. However, this transitional cell type (*DCT-CNT*) is usually not identified or masked by more abundant cell types in single-cell studies. Consequently, most reports identify individual *CNT* and *DCT* clusters^59^. To this end, the kidney atlas can aid the discovery of robust novel markers for *DCT*-*CNT* cells. These markers are detected across the different datasets and can accurately classify *DCT*-*CNT* cells. As demonstrated by the above example, we identify meta-markers for the cell types present in our atlas, including previously known and novel genetic markers. When compared to markers obtained without accounting each individual dataset in an integrated space, meta-markers with a high detection rate can provide replicability that generalizes the cell type identities defined in our atlas. A clear example of such a scenario is *DTL-ATL* cells. As has been described previously, one of the meta-markers identified for this population is *Slc4a11* which expresses a membrane transporter involved in water, ammonia and H^+^ transport. Its expression has been located in DTL cells within the outer medulla and the outer stripes of the inner medulla in mice^46^.

These findings will benefit the broader kidney research community, for example, by aiding the robust *in vivo* identification of cell types. Since human and mouse kidneys show important physiological differences at the cellular level^60^ we believe our work is especially relevant in mouse models. The discovery potential of our atlas, however, is much broader and largely not explored. We acknowledge that, although statistically robust and *in silico* validated with micro-dissected nephron segments^39^, these compendium-wide markers need further *in vivo* validation.

Cellular knowledge of the kidney is likely to change in the coming years. As technologies improve and innovative studies are published, novel cell types will be described. Likewise, cell identities will be re-defined in newer contexts. We aim to incorporate these changes within the atlas in a continuous fashion. We provide a learnt transcriptome-based cell hierarchy that can be easily updated and improved with newer studies, updating the cellular knowledge captured in the compendium. Because we used scVI and scANVI as our integration model, we can leverage scARCHES^61^ to update the latent space of our atlas without retraining. Using this latent space together with the updated scHPL classification tree we can label matching cell types and identify potential novel populations that arise from treatment or disease state. To account for the technical variation of new datasets in the context of the MKA, one can make use of the pre-trained and optimized model we present to obtain an updated latent space that we then use to update the classifier. To make this easily accessible to the community, we share our atlas via a user-friendly web interface, hosted at cellxgene (https://cellxgene.cziscience.com/e/42bb7f78-cef8-4b0d-9bba-50037d64d8c1.cxg/).

In summary, we leverage the large collection of publicly available single-cell and single-nuclei studies and establish a dynamic atlas of the mouse kidney. We demonstrate the extraordinary power of such approach by providing robust markers for elusive cell types. However, the full potential of the created compendium is yet to be explored.

## Acknowledgements

We thank the Chan Zuckerberg Initiative (CZI) for their help with integrating our MKA atlas in cellxgene. We thank investigators for making their data publicly accessible, as it has been essential to this work. We also thank Lieke Michielsen for her help to set up scHPL and useful feedback. This work has received funding from the European Union’s Horizon 2020 research and innovation programme under the Marie Skłodowska-Curie grant agreement No 860977

---

**Figure S1.**
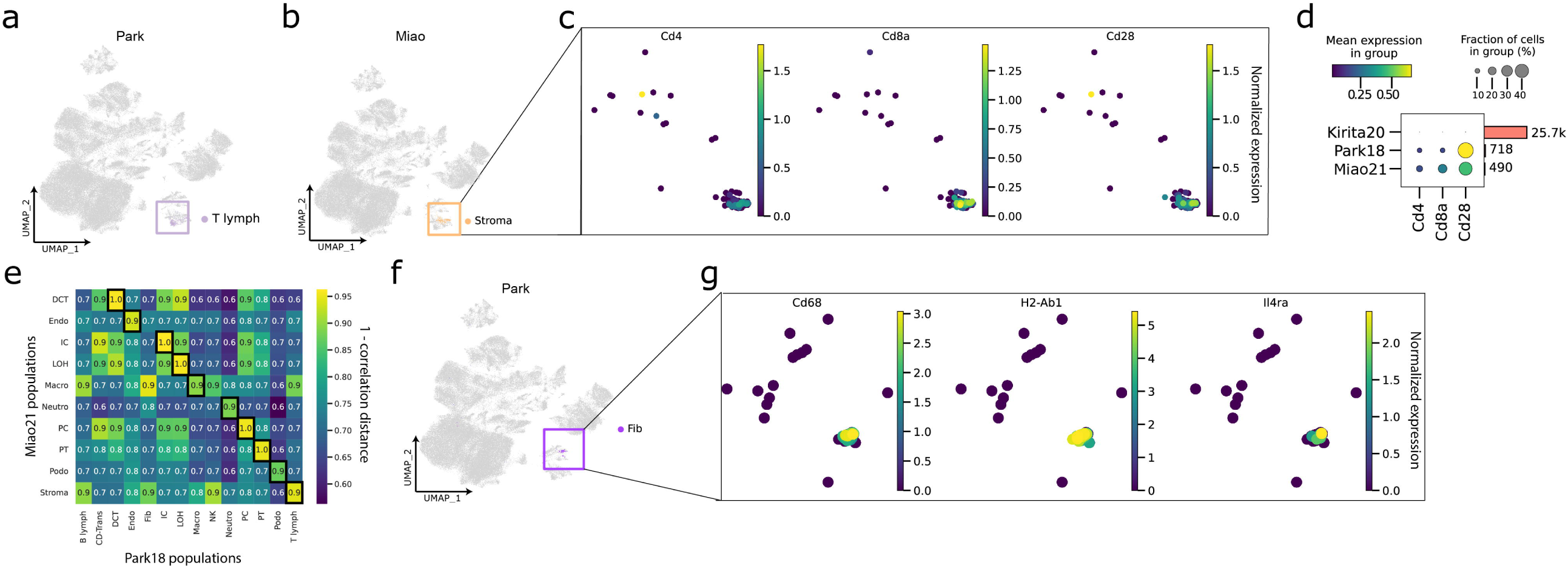
Manual annotation of dissenting cell populations based on scHPL. **(a-b)** UMAP plots, coloured by both *T lymphocytes*_Park18_ (a) and *Stroma*_Miao21_ cells (b). **(c)** UMAP plots of *Stroma* cells coloured by scaled *T lymphocyte* marker expression (*Cd4, Cd8a, Cd28*). **(d)** Dot plot showing the frequency and average level of expression in selected *T lymphocyte* markers (*Cd4, Cd8a, Cd28*) in *Stroma*_Miao21_ cells, *T lymphocytes*_Park18_ as positive control, and Kirita20 non-immune cells as a negative control. The bar plot (right) shows the total cell/nuclei count in each row. **(e)** Heatmap of the similarity (expressed as 1 – correlation distance) of cell populations between Miao21 and Park18 **(f)** UMAP plot, coloured by *Fibroblasts*_Park18_. **(g)** UMAP plots of *Fibroblast* cells coloured by scaled *Macrophage* marker expression (*Cd68, H2-Ab1, Il4ra*).

**Figure S2.**
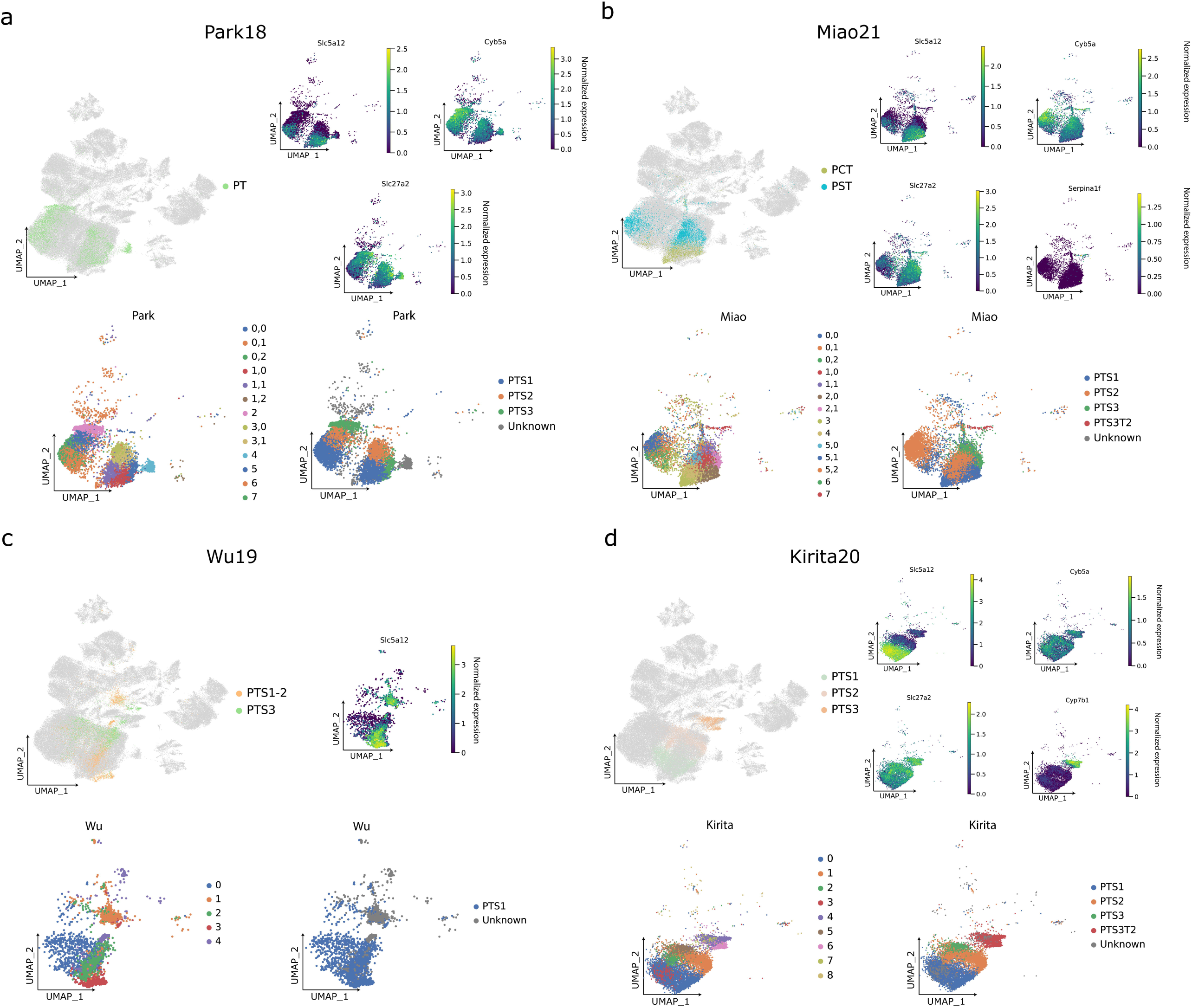
Integrated atlas allows further division of proximal tubule cells. **(a-d)** UMAP plots for Park18 (a), Wu19 (b), Miao21 (c) and Kirita20 (d) highlighting *PT, PTS1, PTS2* and/or *PTS3* (top left), their marker expression (top right; PTS1: *Slc5a12;* PTS2: *Cyb5a;* PTS3: *Slc27a2*, PTS3T2: *Cyp7b1*) and the unsupervised clusters (bottom left). Clusters are renamed to *PTS1, PTS2, PTS3* or *PTS3T2* according to the marker expression overlay (bottom right). If the given marker has no expression on the dataset, the corresponding UMAP plot is omitted. *PT*: Proximal Tubule, *PTS1*: Proximal Tubule Segment 1, *PTS2*: Proximal Tubule Segment 2, *PTS3*: Proximal Tubule Segment 3, *PTS3T2*: Proximal Tubule Segment 3 Type 2.

**Figure S3.**
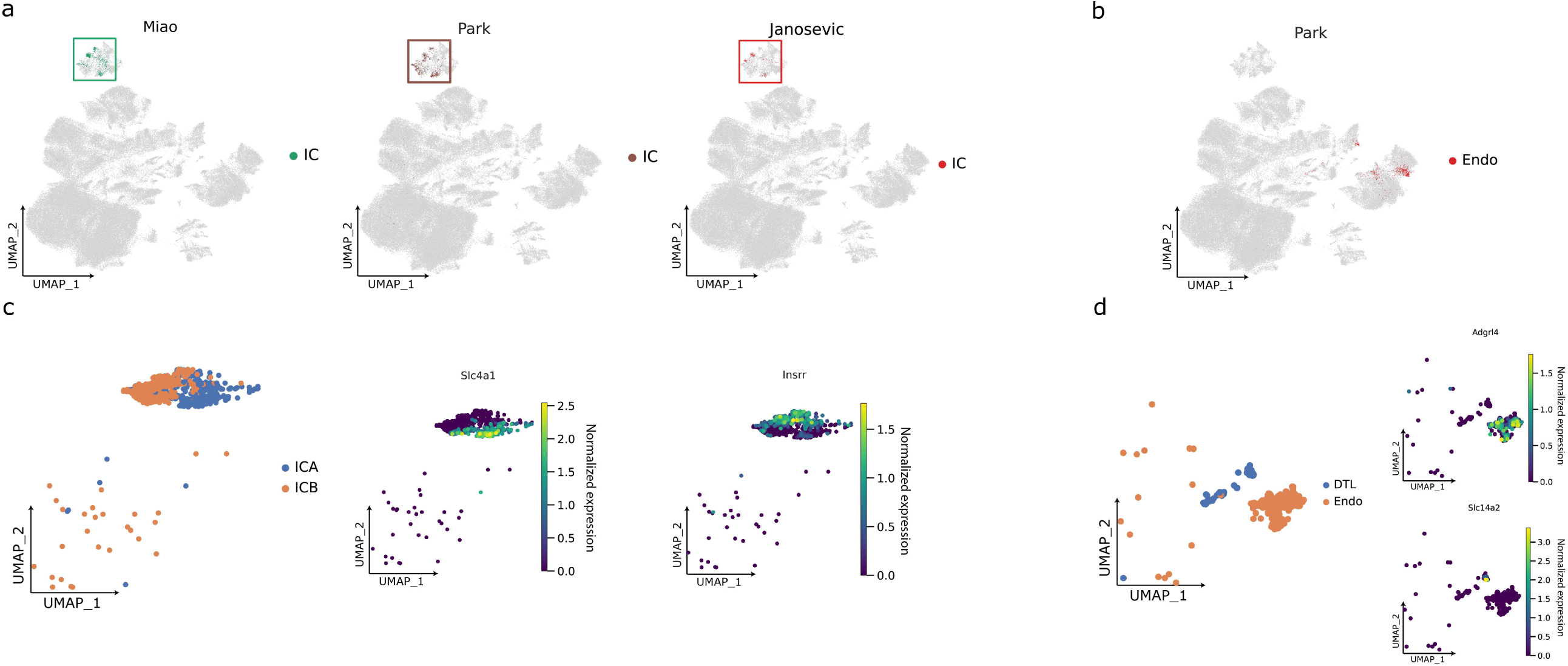
Further division of collecting duct intercalated cells and endothelial cells. **(a-b)** UMAP plots coloured by *IC*_Miao21_ *IC*_Park18_ and *IC*_Janosevic21_ (a) and *Endothelial*_Park18_ cells (b). **(c-d)** UMAP plots coloured by renamed cluster (left) and marker expression (right) of *IC* (c) or *Endothelial* cells (d). Clusters are renamed to *ICA, ICB, Endothelial* or *DTL* according to the marker expression overlay (*ICA*: *Slc4a1*; *ICB*: *Insrr*; *Endo*: *Adgrl4*, DTL: *Slc14a2*).

**Figure S4.**
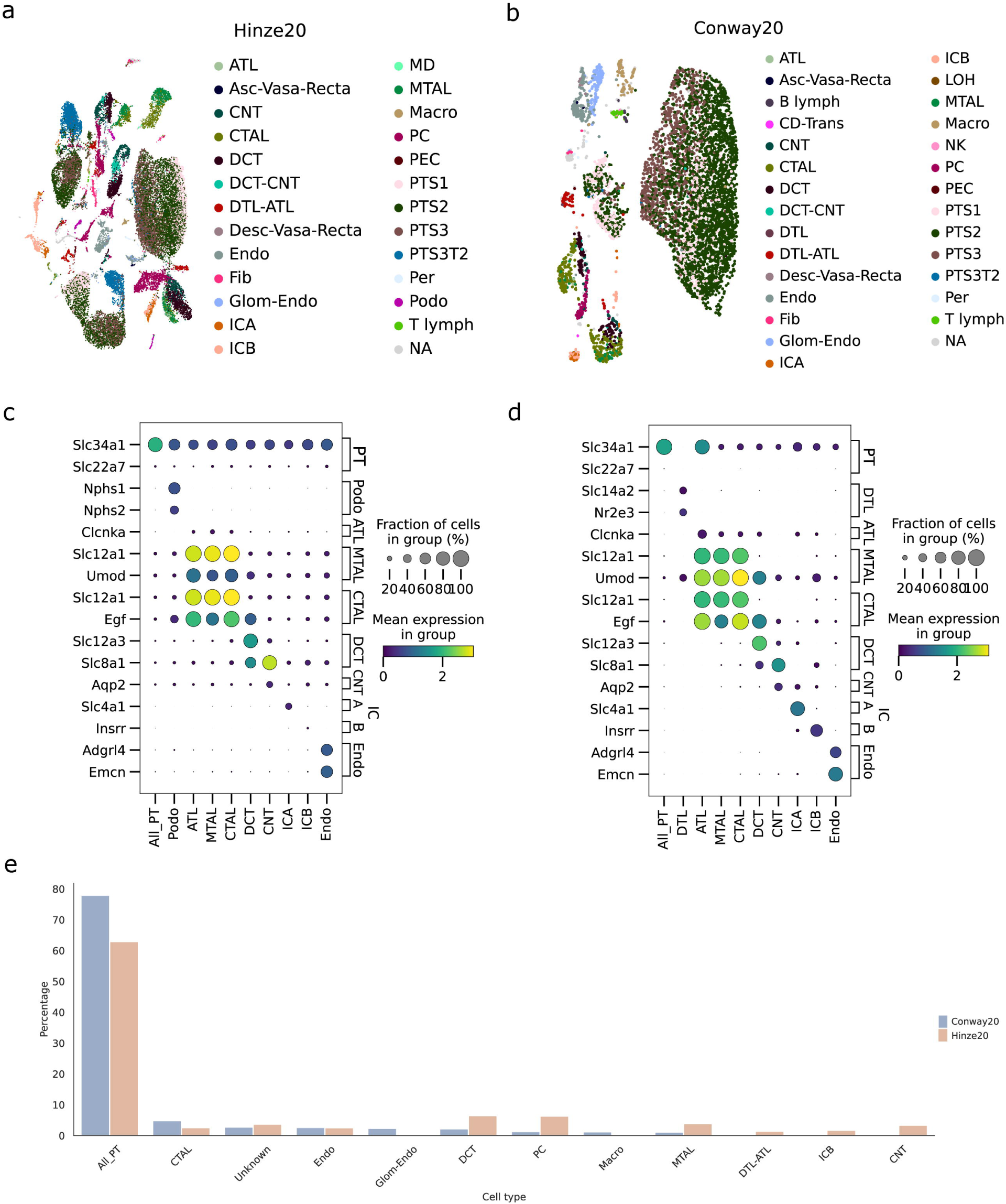
Cell type prediction per dataset. **(a-b)** The final learned hierarchy (classification) tree was used to individually predict the cell types of the unannotated cells in the used datasets in the atlas. **(c-d)** Dotplot of predefined set of standard markers in every predicted population per dataset. *IC*: Intercalated Cell, *ICA*: Intercalated Cell Type A, *ICB*: Intercalated Cell Type B, *Endo*: Endothelial Cell, *Fib*: Fibroblast, *Macro*: Macrophage, *B lymph*: B lymphocyte, *Stroma*: Stroma cell, *NK*: Natural Killer, *T lymph*: T lymphocyte, *PT*: Proximal Tubule, *PTS1*: Proximal Tubule Segment 1, *PTS2*: Proximal Tubule Segment 2, *PTS3*: Proximal Tubule Segment 3, *PC*: Principal Cell, *PEC*: Parietal Epithelial Cell, *Per*: Pericyte, *DCT*: Distal Convoluted Tubule, *ATL*: Ascending Thin Limb of Henle, *MD*: Macula Densa, *LOH*: Loop of Henle, *CTAL*: Thick Ascending Limb of Henle in Cortex, *MTAL*: Thick Ascending Limb of Henle in Medulla, *CNT*: Connecting Tubule, *Podo*: Podocyte, *DTL*: Descending Thin Limb of Henle, *MC*: Mesangial Cell, *Neutro*: Neutrophil, *Asc-Vas-Recta (Asc VR)*: Ascending Vasa Recta, *Desc-Vas-Recta (Desc VR)*: Descending Vasa Recta, *Glom Endo*: Glomeruli Endothelial.

**Figure S5.**
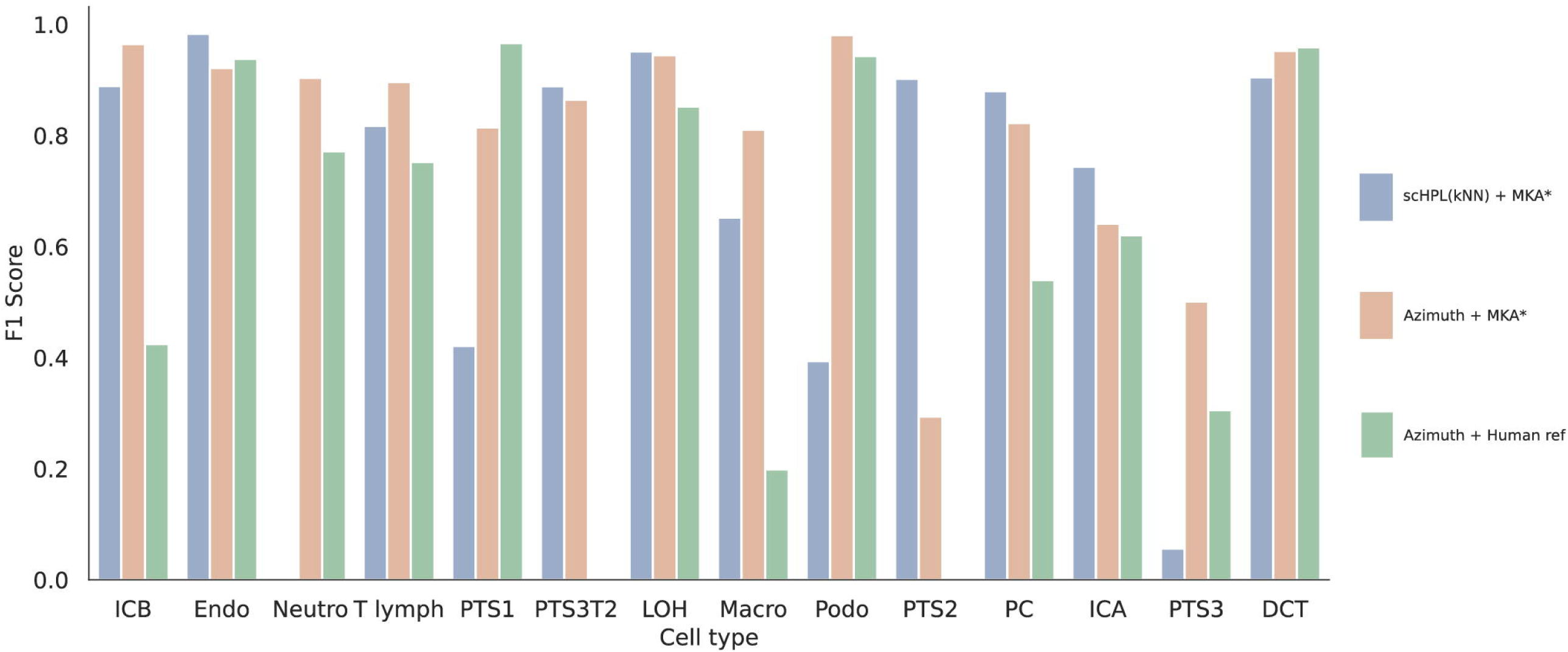
F1 scores per cell type and evaluation experiment. For each cell type, the F1 score is plotted for the different Miao21 classification tasks. Namely, scHPL trained with our partial reference mouse atlas, Azimuth trained with a human reference and Azimuth trained with our partial reference mouse atlas.

**Figure S6.**
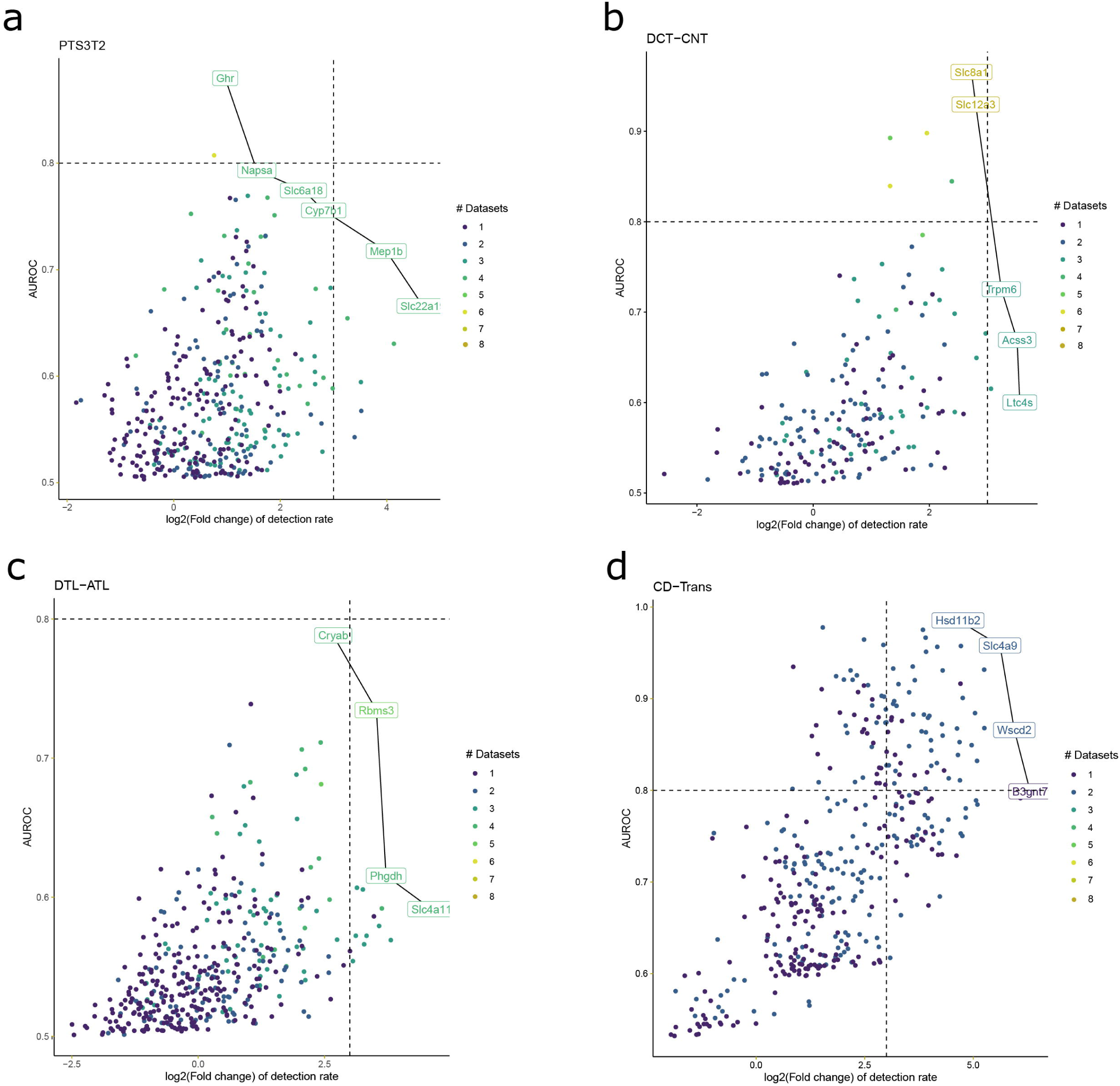
Meta-markers of transitional cell types and rare populations. **(a-d)** Each dot represents a significant meta-marker (FDR < 0.05). Top markers with respect to detection rate (expressed as log2 Fold change) and precision (area under the receiver-operator curve; AUROC) for poorly described cell types are highlighted with a connected boundary line. *PTS3T2*: Proximal Tubule Segment 3 Type 2, *DCT*: Distal Convoluted Tubule, *ATL*: Ascending Thin Limb of Henle, *CNT*: Connecting Tubule, *Podo*: Podocyte, *DTL*: Descending Thin Limb of Henle.

**Table 1. Dataset metadata**. SC: Single-cell, SN: Single-nuclei, QC: Quality control.

**Table 2. Test statistics for the Fisher exact test between the markers defined in the MKA and microdissected nephron segments**. Marker list comparisons are shown per cell type. OR: Odds-Ratio, p: p-value, CI: Confidence interval. *PTS1*: Proximal Tubule Segment 1, *PTS2*: Proximal Tubule Segment 2, *PTS3*: Proximal Tubule Segment 3, *DCT*: Distal Convoluted Tubule, *ATL*: Ascending Thin Limb of Henle, *CTAL*: Thick Ascending Limb of Henle in Cortex, *MTAL*: Thick Ascending Limb of Henle in Medulla, *CNT*: Connecting Tubule, *DTL*: Descending Thin Limb of Henle.

**File name: Supplementary Data 1**

**Description:** Differentially expressed genes for every cell type compared to all other cells (related to Fig. 5A) in the MKA. Scores: z-score underlying the computation of the p-value for each gene for each group. Log2FC: log2 fold change for each gene for each group. P-values were adjusted for multiple testing using the Benjamini-Hochberg correction method. This list was obtained using a Wilcoxon rank-sum test, as described in Materials and Methods.

**File name: Supplementary Data 2**

**Description:** List of identified meta-markers per cell type together with AUROC and fold change of detection rate obtained from the MKA as described in Materials and Methods (related to Figs. 5A and 5B). Genes are ranked by recurrence (number of datasets in which the gene is differentially expressed) and by AUROC.

